# Pangenomic structural variant patterns reflect evolutionary diversification in *Brassica napus*

**DOI:** 10.1101/2024.12.19.629214

**Authors:** Nazanin P. Afsharyan, Agnieszka A. Golicz, Rod J. Snowdon

## Abstract

**Background:** Understanding genetic diversity is crucial for enhancing crop productivity. This study explores species-wide genome structural variation and its role in intraspecific and ecogeographical diversification of *Brassica napus*, a recently evolved, globally important allopolyploid crop.

**Results:** We performed whole-genome long-read DNA-sequencing and constructed reference-guided genome assemblies for 94 diverse, homozygous accessions, including winter-type, spring-type, and East Asian oilseed, along with kale forms and swedes/rutabagas. We investigated pangenomic patterns of genomic structural variants (SV) and determined pangenome-wide distributions and frequencies of inversions, gene presence-absence variants, and collective SV including insertions and deletions. Results revealed pangenome-wide patterns for insertions, deletions, inversions, and large chromosomal deletions/duplications, reflecting evolutionary diversification across morphotypes and ecotypes. Collective SV were unevenly distributed and biased toward subgenome A, with asymmetrical selection pattern favoring subgenome C. Selection signatures for inversions exhibited no subgenome asymmetry; however, selection signal strength and frequency increased in paracentric chromosome regions, highlighting their evolutionary significance. Selective sweep analysis identified regions for collective SV and inversions, harboring genes for organ formation, cell division and expansion in swede, and stress responses in East Asian oilseed rape. Large chromosomal duplications and deletions distinguished swede from oilseed rape, particularly in subgenome C, including copy-number variation in flowering-time genes *BnFLC.C09* and *BnATX2.C08*, and cell wall development gene *BnCEL2.C08*.

**Conclusions:** These findings underscore functional and evolutionary significance of pangenomic SV formation during *B. napus* diversification. Information on SV patterns with putative functional relevance, provides breeding insights, particularly for developing molecular markers to optimize performance of *B. napus* and other *Brassica* crops.

## Background

Genomic structural variants (SV) are abundant chromosomal or genic modifications, typically at least 30 base pairs (bp) in size. SV include deletions, insertions, duplications, inversions, translocations, chromosomal losses and gains, and complex DNA rearrangements [1,2]. Such variants lead to segmental gene presence-absence variation (PAV) and copy-number variation (CNV). In polyploids with multiple similar subgenomes, chromosome-scale homoeologous exchanges are also frequent [3–5]. All of these events can impact genes, and hence, influence all kinds of agronomic traits in crops [6]. SV events have been shown to play a crucial role in adaptation and domestication of crops, as they affect gene structure and genomic regulatory regions more significantly than smaller variants such as single nucleotide polymorphisms (SNPs) and small insertions or deletions (indels). Several mechanisms contribute to SV formation, including the insertion of transposable elements (TEs), double-strand break repair, recombination errors, or meiotic crossovers among non-homologous chromosomes [2].

Polyploidy in plants can result from spontaneous genome duplication (autopolyploidy) or interspecific hybridization (allopolyploidy). Many recent polyploid crops have undergone multiple rounds of polyploidization [7] or have been subjected to introgressive hybridization with gene pools of different ploidy levels [8,9], enabling them to adapt to new environments [10]. Plant genomes, characterized by high levels of repetitive sequences and varying degrees of polyploidy, add complexity to the relationship between SV and the occurrence of crossovers during meiosis. For example, recent studies have shown that large SV, particularly inversions, can suppress recombination. However, crossovers within these SV can sometimes occur, and depending on their relative position to the centromere, these may result in unbalanced gametes with dicentric and/or acentric chromosomes, or balanced gametes with new duplications and deletions [11].

The major crop species *Brassica napus* L. (genome AACC, 2n = 38) is estimated to have emerged no later than around 7000 years ago [3,12,13] from the spontaneous hybridization of its diploid progenitors *B. rapa* and *B. oleracea*. Because no wild *B. napus* forms are known, the actual hybridization event likely did not occur before the early middle ages, when the two progenitor species from Asia and the Mediterranean region are thought to have been cultivated together for the first time as a result of human trade and exchange along the Silk Road. Subspecies *B. napus ssp. napus* is today a globally important oilseed crop (rapeseed, oilseed rape, canola) which also includes leafy kale or fodder forms, while the other subspecies *B. napus ssp. napobrassica*, known as swede or rutabaga, has an enlarged hypocotyl that is consumed as a vegetable or used for livestock grazing [14]. Despite a short domestication history, *B. napus* exhibits remarkable phenotypic diversity and phenological adaptation. Recent genomic studies have revealed extensive chromosome-scale SV events contributing to this diversity. However, these studies also highlight the challenges of detecting whole-genome SV in the allotetraploid genome of *B. napus* due to high homoeology between chromosomes and frequent homoeologous exchanges [3]. These challenges have been mitigated by recent advances in long-read sequencing technologies and bioinformatics tools, which allow for high-precision SV detection based on long-read alignments [15,16] or accurate whole-genome assemblies [17].

*B. napus* has adapted to diverse climate zones in major production areas across Europe, Asia, North and South America and Australia, requiring broad diversity in adaptation traits including growth habit, vernalization requirement, winter hardiness and phenology, from early-flowering spring types to late-flowering winter and biennial types. This makes it a valuable model for studying intraspecific diversification and ecogeographic adaptation [3]. There are three major oilseed *B. napus* ecotypes. Winter-type oilseed rape (WOSR) originated in Europe and is sown in Autumn, being adapted to cold winters with a stringent vernalization requirement and a longer life cycle [13]. Spring-sown oilseed rape (SOSR), known particularly in North America as canola, was domesticated in Europe around 1700 and has no vernalization requirement [18]. An intermediate form, semi-winter type oilseed rape, was developed in East Asia over the past 200 years for adaptation to milder winters [19]. In breeding efforts to improve local adaptation and disease resistance of semi-winter type oilseed rape, interspecific crosses with diploid progenitors, particularly the subgenome A donor *B. rapa,* have been used [20].

The origin of *B. napus*, including swede, remains debated with recent studies suggesting a single hybridization event [21], while others suggest a few independent hybridization events [22]. Swede is thought to have been domesticated before oilseed rape (OSR) in Europe, and gene introgression from *B. rapa* into swede [21] has been used particularly to transfer resistance to soil pathogens [23]. Major breeding breakthroughs in the 1960s and 1970s eliminated undesirable erucic acid in the oil and reduced aliphatic glucosinolates in the seed meal, drastically improving the value of oilseed forms as a high-quality source of vegetable oil and protein-rich livestock feed. Since then, intensive breeding in OSR has considerably improved oil content, seed yield, and disease resistance. However, these breeding efforts have also narrowed the gene pool of elite cultivars, particularly because only a few trait donors were utilized to establish the key seed quality traits [24].

Previous studies have investigated species-wide genetic variation, including SNPs [3,13,21,25,26], simple sequence repeats (SSRs) [27], copy number variations (CNVs) [28], and homoeologous chromosomal exchanges [29] in *B. napus* to study its origins and ecogeographical adaptation. Several of these studies used the global *B. napus* ERANET-ASSYST collection [27] to investigate genome-wide genetic variation using SSR, CNV and SNP variants [27,28,30]. Recently, there has been growing interest in constructing pangenome references to describe the available genetic diversity within crop species. Compared to single-reference genomes, pangenome references enable more comprehensive and accurate identification and representation of genetic variants within a species [31]. Advances in high-throughput, cost-effective DNA sequencing technologies have led to a significant increase in the development of pangenome references for major crops, including rice (*Oryza sativa* L.) [32,33], maize (*Zea mays* L.) [34], soybean (*Glycine max* L.) [35], and rapeseed [36,37]. Recent publications have also introduced pangenome SV catalogues for *Arabidopsis* [38] and a pangenome inversion index for rice [39]. Despite the growing body of research on functional variation caused by SV, and the increasing precision and affordability of third-generation sequencing technologies, no large-scale study has yet surveyed the species-wide SV landscape in *B. napus*, or its connection to intraspecific diversification.

In this study, we analyzed 94 homozygous, ecogeographically diverse accessions of *B. napus* from the ERANET-ASSYST diversity collection to explore the pangenomic SV landscape and its relationship to intraspecific diversification. Based on alignment to the gold-standard *B. napus* Darmor-bzh v10 reference genome [40], we described and characterized pangenome-wide distribution and frequencies for gene presence-absence variants, inversions, and collective SV. We performed long-read DNA sequencing using Oxford Nanopore Technology PromethION R9.4.1 flow cells, achieving an average 39.48× whole-genome coverage. This data was used to construct reference-guided genome assemblies. SV calling and inversion detection were performed using two approaches: long-read alignments and genome assembly alignments. We conducted genome-wide analyses of phylogeny, selective sweep signatures, and presence-absence variation between different ecotypes and morphotypes. Additionally, we investigated large chromosomal deletions and duplications, and analyzed their occurrence in different morphotypes.

## Results

### Generating high-coverage genome-wide long-read DNA sequencing data

Using Oxford Nanopore technology, we performed long-read whole-genome DNA sequencing of 94 *B. napus* accessions (Additional file 1: Table S1) that span the range of genetic diversity represented by the ERANET-ASSYST diversity set [27]. Overall, we generated approximately 39 Tb of Fast5 data. Given the *B. napus* Darmor-bzh v10 (D10) reference genome size of 1,132 Mbs [40], coverage ranged between 21.63× and 71.52× (24.49 Gb to 80.97 Gb of fastq data per accession), with an average coverage of 39.48×. N50 values ranged from 18.00 kbp to 54.61 kbp, with an average of 29.37 kbp (Additional file 1: Table S2).

### Constructing alignment-based genome assemblies

We constructed genome assemblies for all 94 *B. napus* accessions by building de novo contigs using long reads, followed by alignment-based scaffolding based on the D10 reference assembly. The average assembly size was 0.89 Gb across 19 chromosomes, with individual assembly sizes ranging from 1.01 Gb to 0.66 Gb. The assemblies were evaluated using Benchmarking Universal Single-Copy Orthologs (BUSCO), revealing that on average, 99.57% of 1,614 complete BUSCO genes were detected across the 94 assemblies (Additional file 1: Table S3). Of these, the average percentage of complete single-copy and duplicated BUSCOs was 5.48% and 94.09%, respectively, reflecting the polyploid nature of *B. napus*.

### Determining frequency of collective SV demonstrated pangenome-wide SV distribution in intergenic and intragenic regions

SV including insertions, deletions, inversions, and translocations (denoted here as collective SV) were detected after aligning long reads to the D10 reference genome. The mapping efficiency of the filtered reads was 100%. Total 69,004 SV from the call set overlapped with large deleted chromosome segments and were filtered-out as summarized in Additional file 1: Table S4 and S5. The SV detection pipeline identified a grand total of 3,649,887 quality-filtered SV across all accessions, consisting of 1,922,316 insertions, 1,727,416 deletions, 139 inversions and 16 translocations (Fig. 1a, Additional file 1: Tables S6 and S7). In total, pangenome-wide frequencies of 861,307 SV events, up to 100 kbp in size, were determined based on long-read alignments including 755,613 insertions, 105,655 deletions, 23 inversions, and 16 translocations (Additional file 1: Table S8). Of 108,190 genes annotated in the D10 reference genome, 39.58% were found to contain SV in their coding sequences (CDS) or untranslated regions (UTRs), including rare variants (Additional file 1: Table S9). Among these were 108 putative flowering-time-related genes (Additional file 1: Table S10) and 174 putative resistance (R) genes (Additional file 1: Table S11). Comparisons with previously reported SV confirmed, for instance, a 288 bp SV in the *FLOWERING LOCUS T* gene on chromosome A02, as identified by Vollrath et al. [41] through both short-read sequencing and PCR validation. Additionally, we confirmed a 1.3 kbp SV in the putative promoter region of the same gene reported by Chawla et al. [42].

**Figure 1:**
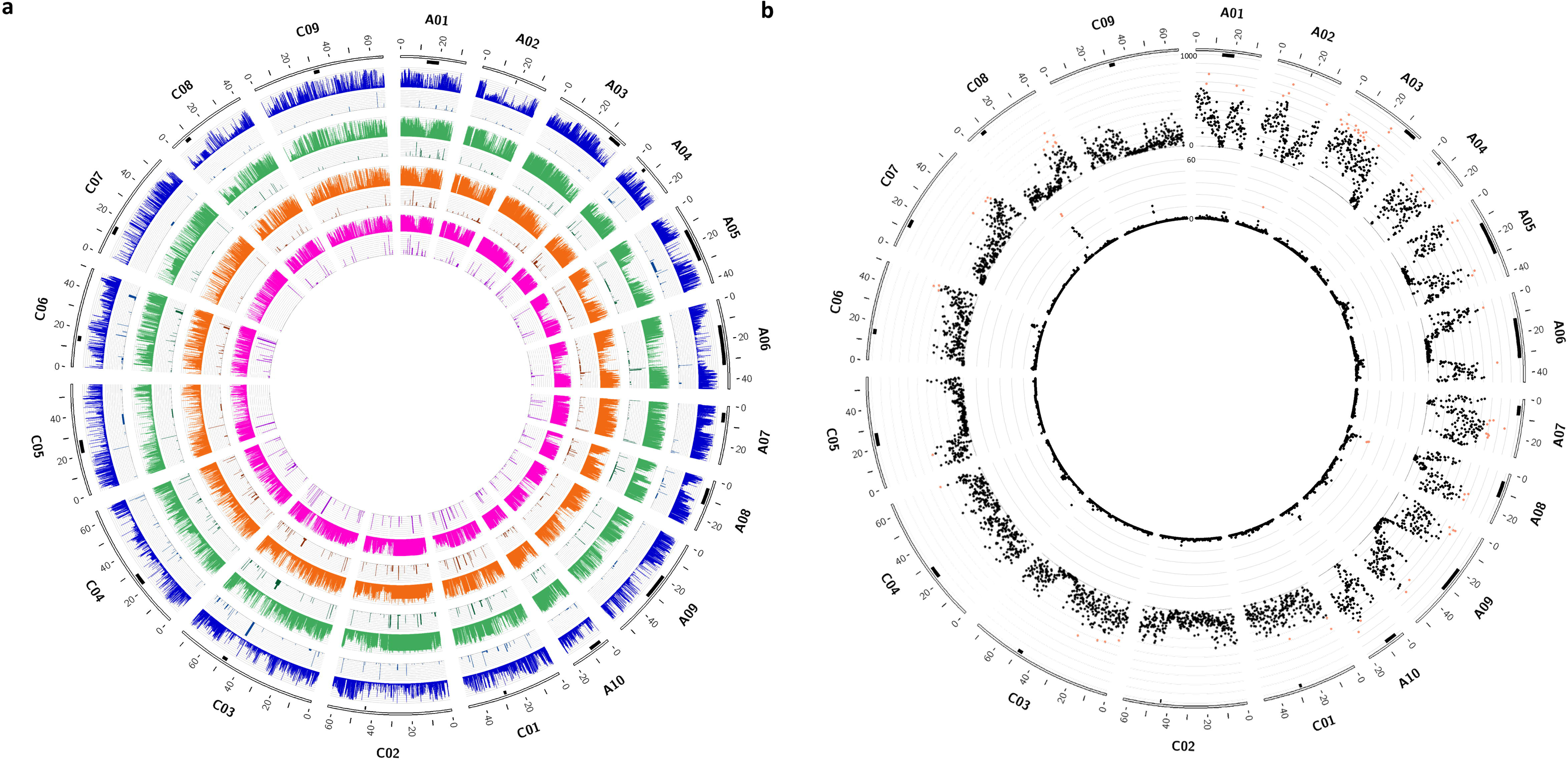
Overview of pangenome-wide distribution and frequency of collective SV (Pan-CSV) and inversions (Pan-INV) in a panel of 94 homozygous, ecogeographically diverse accessions of *B. napus*. a) The distribution (x-axis) and frequency (y-axis, 0 to 100) of SV events for the pan-CSV and pan-INV. For each ecotype group, the histogram on top describes pan-CSV, while the histogram on the bottom shows pan-INV. The circos plot from outer to inner layer includes chromosomes, centromeric regions (marked in black), WOSR (blue), SOSR (green), East Asian OSR (orange), and swede/rutabaga (magenta) accessions. The frequency is normalized as a percentage. b) Chromosome distribution (x-axis) and number (y-axis, outer plot: 0 to 1000, inner plot: 0 to 60) of hotspots for pan-CSV (outer plot) and pan-INV (inner plot). The chromosomes are shown as the outer layer, followed by centromeric regions (marked in black). The dot plots display the number per 200 kbp window. The red dots indicate the top 30% per 200 kbp window, calculated separately for each subgenome. The coordinates of the centromeric regions were determined as detailed by Rousseau-Gueutin et al. [40]. Plots were generated using Circos [91].

Furthermore, SV in the *FLOWERING LOCUS C* gene on chromosome A10 included a 621 bp SV and a 4,421 bp SV in the promoter region, and a 5,565 bp SV in the first exon, all previously reported by Song et al. [43].

### Determining pangenome-wide distribution and frequency of inversions revealed link with intraspecific diversification

Reference-guided genome assemblies were aligned to the D10 reference genome to detect 1,166 non-redundant inversion events, ranging in size from 199 bp to 12.097 Mbp, including 84 inversions larger than 1 Mbp (Fig. 1a, Additional file 1: Tables S12 and S13). These inversions were merged, and their pangenome-wide distribution and frequency was determined. Analysis of over-represented inversions in each ecotype group revealed 1, 8, and 15 inversions present in ≥70% of the accessions in the WOSR, SOSR, and swede/rutabaga groups, respectively. Among them a 506.4 kbp inversion was located on chromosome C05 which was present in all swede accessions but was not found in WOSR and SOSR ecotypes. On the other hand, there were 5 and 1 inversions that were under-represented (≤30%) in WOSR and SOSR respectively (Additional file 1: Table S14).

### Hotspots for pangenome-wide collective SV and inversions

We investigated SV hotspots in 200 kbp windows across all chromosomes for pangenome-wide distribution and frequencies of collective SV and inversions. For subgenomes A and C, an average of 294.43 (SD = 189.23) and 131.67 (SD = 100.44) SV events per 200 kbp were detected for collective SV, while for inversions, an average of 0.87 (SD = 1.99) and 0.44 (SD = 2.41) events were found for each subgenome, respectively. Statistical analysis showed significantly higher SV occurrence in subgenome A compared to subgenome C for both collective SV (*p*-value = 1.68E-191) and inversions (*p*-value = 7.76E-11). Hotspots were defined as the top 30% of windows with the highest frequency of SV. For the collective SV, 55 hotspots were found in subgenome A and 23 in subgenome C, mostly towards telomeric regions (Fig. 1b, Additional file 1: Table S15). For the inversions, hotspots were found in chromosomes A02 and C08, near centromeres, and A08, within centromeric regions (Fig. 1b, Additional file 1: Table S16).

### Structural variants differentiate accessions by ecotype and subspecies

Unrooted phylogenetic analyses of the subgenomes A and C based on pangenome-wide collective SV and inversions clustered most accessions into their respective WOSR, SOSR and swede groups. As expected, East Asian (generally semi-winter) OSR accessions were intermediate. The A lineage analysis grouped swedes and some East Asian OSR accessions closer to *B. rapa* for both collective SV and inversions, while the C lineage analysis clustered some East Asian OSR accessions near wild *B. oleracea* for collective SV (Fig. 2b) and grouped wild *B. oleracea* closer to SOSR and East Asian OSR accessions for inversions (Fig. 2c). Principal component analysis (PCA) supported these findings, clearly separating WOSR, SOSR, and swede groups for both collective SV and inversions (Fig. 2d).

**Figure 2:**
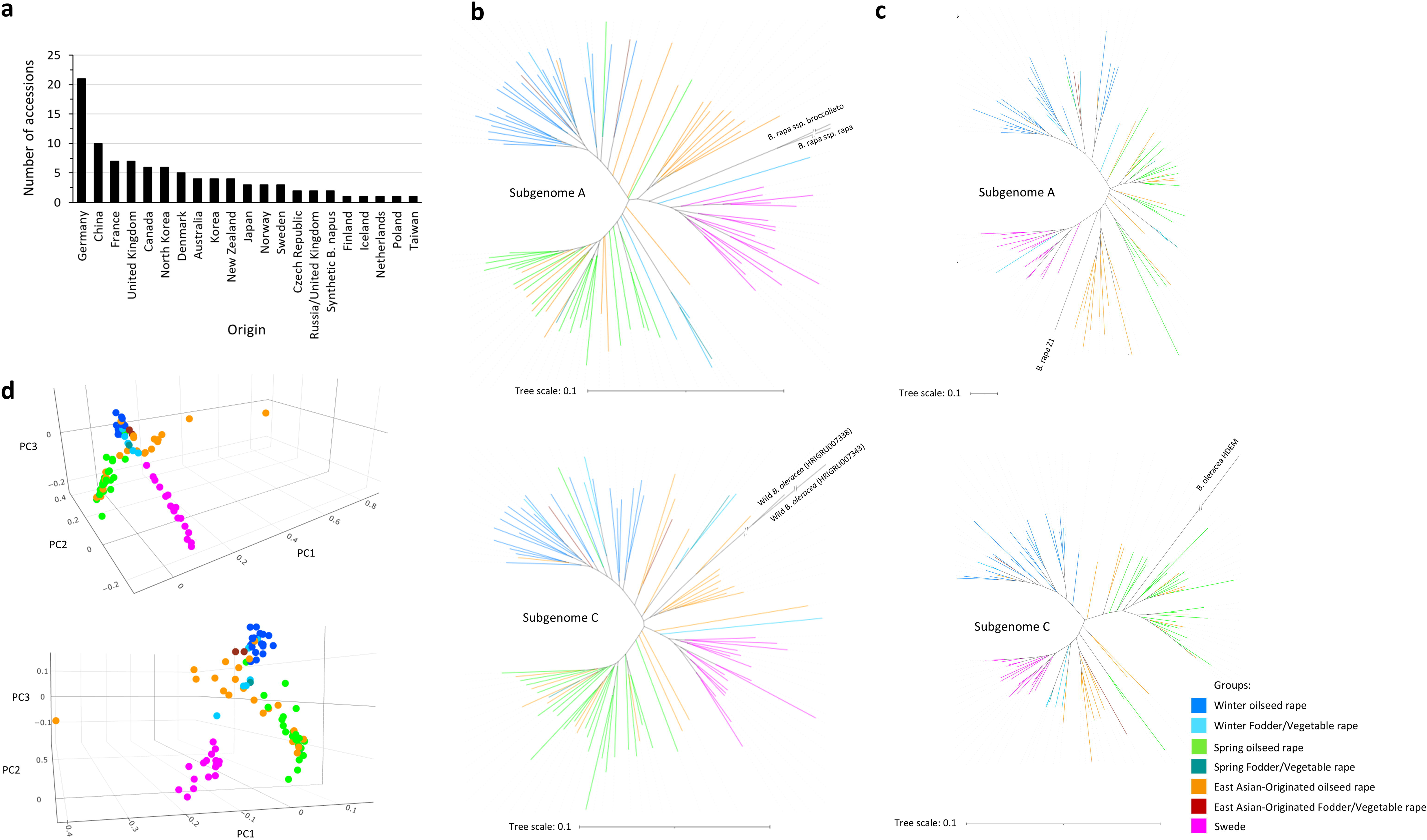
Phylogenetic analysis of 94 homozygous, ecogeographically diverse accessions of *B. napus* based on pangenome-wide collective SV and inversions. a) Overview of the origin and number of accessions. b) Phylogenetic tree of A and C lineages for all accessions, constructed based on pangenome-wide distribution and frequency of collective SV (pan-CSV). Two accessions of *B. rapa* (*B. rapa* ssp. *rapa* and *B. rapa* ssp. *broccolieto*) [61] and two accessions of wild *B. oleracea* (Genebank/cultivar ID: HRIGRU007343 and HRIGRU007338) [62] were included in the analysis for the A and C lineages, respectively. c) Phylogenetic tree of A and C lineages for all accessions, constructed based on pangenome-wide distribution and frequency of inversions (pan-INV). *B. rapa* accession Z1 and *B. oleracea* accession HDEM [63] were included in the analysis for A and C lineages, respectively. d) PCA plot of all accessions based on pan-CSV (top) and pan-INV (bottom). Phylogenetic trees are constructed using a maximum likelihood approach. The reliability of the tree was also confirmed by 1000 bootstrap replicates, and visualized using the Interactive Tree Of Life online tool (https://itol.embl.de).

### Selective sweep signatures based on pangenome-wide collective SV exhibited subgenome bias for intraspecific diversification

Using the pangenome-wide distribution and frequency of collective SV, we investigated selection signatures contributing to diversification in *B. napus*, identifying key chromosome regions in pairwise comparisons of swede vs. non-swede, East Asian OSR vs. all other OSR, and WOSR vs. SOSR accessions (Fig. 3a; Additional file 1: Tables S17-19). Then, we determined the genes that were located in regions under selection (Additional file 1: Tables S20-22). Significantly higher average z-values for subgenome C were observed compared to A, for swede vs. non-swede (*p*-value = 5.05E-11) and WOSR vs. SOSR (*p*-value = 1.04-09) pairwise comparisons. No significant differences in average z-values were noted for East Asian OSR vs. all other OSR accessions. GO enrichment analysis for swede vs. non-swede accessions revealed significant overrepresentation of genes involved in cell division and expansion, meiosis, and environmental responses and disease resistance (Additional file 1: Tables S23). For East Asian OSR vs. all other OSR groups, regions under selection showed significant overrepresentation of genes involved in disease resistance, and abiotic stress response (Additional file 1: Tables S24). Similarity, enriched GO terms in genomic regions under selection for WOSR vs. SOSR indicated presence of genes potentially involved in disease resistance and stress response (Additional file 1: Tables S25). Additionally, the key flowering time genes, orthologs of Arabidopsis *FLOWERING LOCUS C* (*FLC*), in chromosomes A03 and A10 were also located near genomic regions with significant selective sweep signatures for WOSR vs SOSR comparison. Since *FLC* is a key determinant of vernalization requirement, it can be expected that SV within *B. napus* orthologs of this and other key adaptive genes might indeed have contributed to intraspecific selection in *B. napus*.

**Figure 3:**
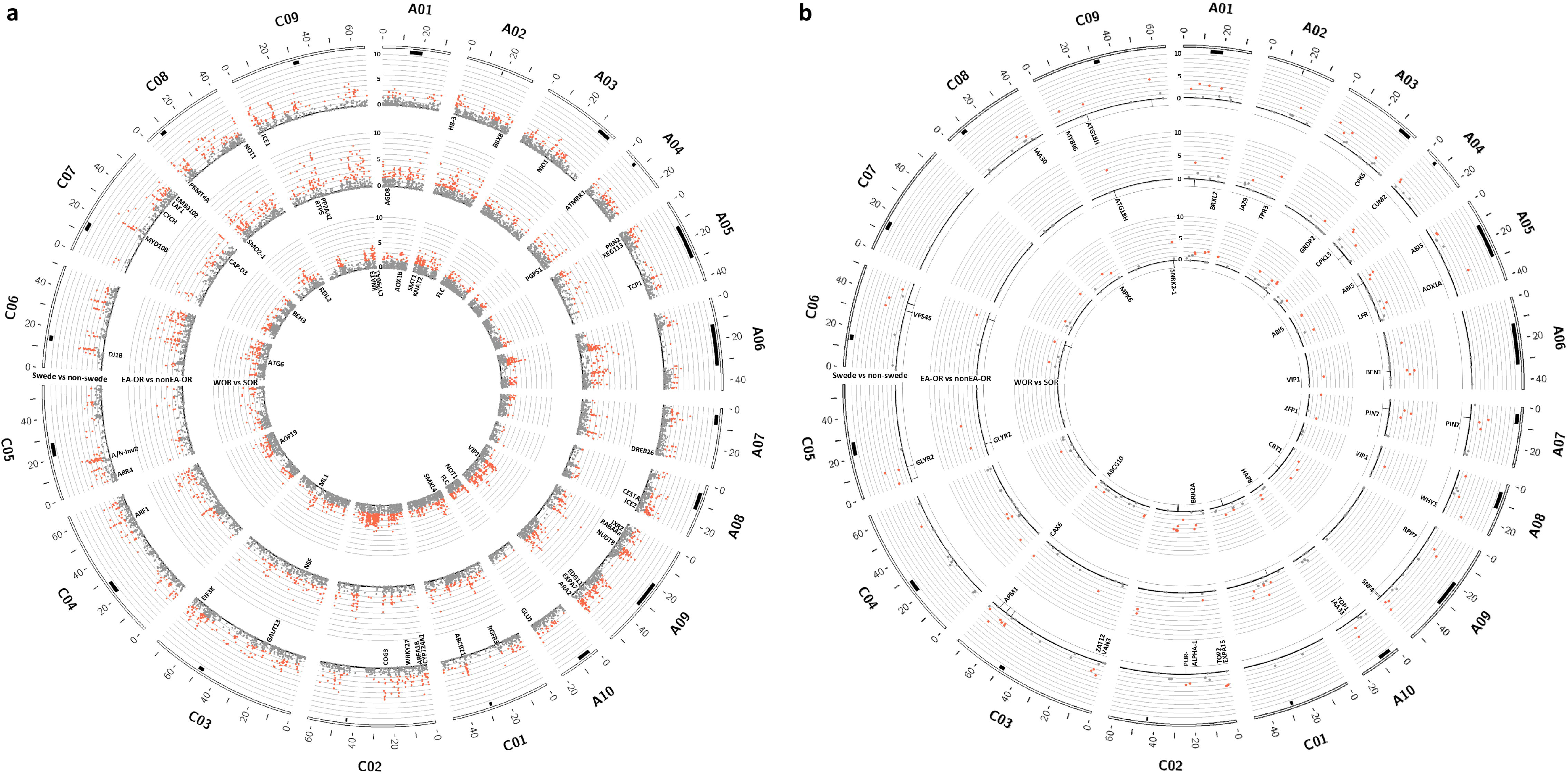
Overview of pangenome-wide screening for selective sweep signatures associated with intraspecific and ecogeographical diversification in *B. napus*. a) Distribution of pangenome-wide selective sweep signals for collective SV (pan-CSV). b) Distribution of pangenome-wide selective sweep signals for inversions (pan-INV). Positions of the overlapping signals from collective SV and inversions are marked below each Manhattan subplot in Figure b. The chromosomes are shown as the outer layer, followed by centromeric regions (marked in black). The Manhattan plots, from outermost circle inwards, depict FST values for pairwise comparisons of swede vs. non-swede accessions, East Asian OSR vs. all other OSR accessions, and WOSR vs. SOSR accessions, respectively. A 50 kbp non-overlapping window was used to calculate FST values. FST values were normalized as z-scores. Dots colored in red indicate values above the significance level threshold of α = 0.05, corresponding to z = 1.645. A number of genes located within the regions under selection, known to be related to developmental and adaptation mechanisms, are presented under the respective Manhattan plot. The coordinates of the centromeric regions were determined as detailed by Rousseau-Gueutin et al. [40]. Plots were generated using Circos [91].

### Selective sweeps based on pangenome-wide inversions highlighted stronger signals towards paracentric regions

The pangenome-wide distribution and frequency of inversions was used to explore selection signatures during domestication and breeding of *B. napus*. The results revealed chromosome regions which contribute to diversification in all pairwise comparisons, including swede vs. non-swede, East Asian OSR vs. all other OSR, and WOSR vs. SOSR accessions (Fig. 3b; Additional file 1: Tables S26-28). A comparison of average z-values for subgenomes A and C found no significant differences. The detected genomic regions under selection were annotated to determine genes located in those regions (Additional file 1: Tables S29-31). GO enrichment analysis for swede vs. non-swede accessions revealed significant overrepresentation of genes involved in cell division and expansion, environmental responses, and disease resistance (Additional file 1: Tables S32). Analysis for East Asian OSR and all other OSR accessions pairwise comparison, found significant enrichment for various genetic pathways including genes related to development, environment interaction, and abiotic stress responses (Additional file 1: Tables S33). GO enrichment analysis for comparison of WOSR vs. SOSR accessions, detected enrichment for genes involved in reproductive growth and development, environmental interaction, and abiotic stress response in regions under selection (Additional file 1: Tables S34). Since some inversions may be in linkage disequilibrium (LD) with insertions and deletions, or contain them, we also investigated the regions under selective sweeps that overlapped with the selective sweep signatures identified for collective SV. A total of 14, 9, and 7 selective sweep regions overlapped with those detected using collective SV for the pairwise comparisons of swede vs. non-swede, East Asian OSR vs. all other OSR, and WOSR vs. SOSR accessions, respectively (Fig. 3b; Additional file 1: Tables S35-37). These findings suggest that the inversions have contributed considerably to intraspecific selective signatures resulting from intraspecific morphotype diversification and human selection in *B. napus*.

### Pangenome-wide gene PAV revealed association with morphological and intraspecific diversification

Genomic rearrangements, including large duplicated or deleted genomic segments, give rise to large-scale gene presence-absence variation. Analyzing the depth of reads aligned to the D10 reference genome for each accession revealed genomic segments of ≥10 kbp that were either duplicated or deleted. The chromosomes of subgenome C exhibited a higher number of large duplicated and deleted regions compared to those of subgenome A, with segmental deletions being more frequent than duplications. In total, 103,163 genes were affected by these large duplicated or deleted segments across the entire panel (Additional file 1: Table S38). From these, 829 genes were over-represented among either WOSR, SOSR, East Asian OSR or swede accessions (≥70% in the target group and ≤30% in the other groups). Notably, in all swede accessions, large segmental deletions were observed on chromosomes C01, C02, C08, and C09, while segmental duplications were found on chromosome A09. The East Asian OSR and WOSR groups exhibited small deletions in all accessions on chromosomes C03 and C05, respectively (Fig. 4; Additional file 1: Table S39). Comparative analysis between the swede and non-swede groups highlighted genes located in duplicated and deleted regions, some showing more than one copy number change, that are potentially involved in cell wall development and expansion, lignin biosynthesis, and flowering. Specifically, large deleted segments in subgenome C, present in all swede accessions, contained one copy of *BnCEL2.C08* and *BnEXPA7.C08*, and two copies of *BnATX2.C08* and *BnFLC.C09*, which are orthologs of *Arabidopsis CELLULASE 2 (CEL2)*, *EXPANSIN A7 (EXPA7)*, *TRITHORAX 2 SET DOMAIN PROTEIN 30 (ATX2)*, and *FLOWERING LOCUS C (FLC)*, respectively. Additionally, large duplicated regions in most swede accessions carried two copies of *BnCEL2.A09* but just one copy each of *EXPA7.A09*, *BnATX2.A09*, and *BnFLC.A10*. These findings demonstrate that large chromosomal deletions and duplications are associated with the diversification of swede and non-swede groups, further supporting the role of large structural variants in the intraspecific diversification of *B. napus*.

**Figure 4:**
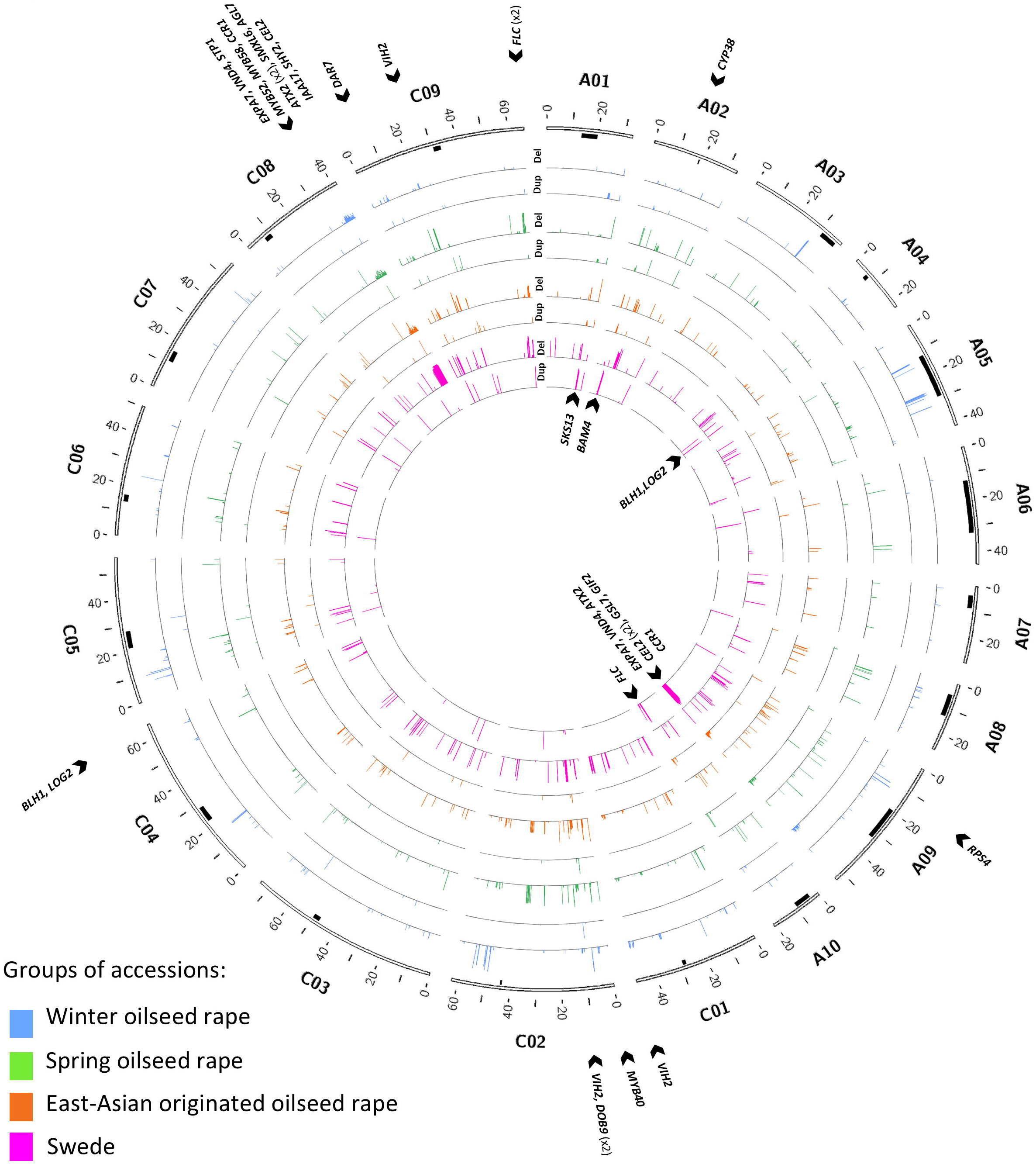
Overview of pangenome-wide gene presence-absence variations correlating with ecotype diversification in diverse accessions of *B. napus*. From outermost circles inwards, each pair of plots describes results for WOSR (blue), SOSR (green), East Asian OSR (orange), and swede/rutabaga (magenta) accessions. The distribution (x-axis) and percentage (y-axis, 0 to 100) of genes whose deletion or duplication correlated with each ecotype group are presented based on their presence in ≥70% of the target ecotype group and ≤30% of other ecotype groups. EA-OR accessions were not included as “other ecotypes,” as some accessions of this group clustered closely with WOSR and SOSR accessions in principal component analysis and phylogenetic analysis. The candidate genes that may correspond to ecotype diversification in deleted and duplicated regions are indicated outside and inside the circos plot, respectively. Plots were generated using Circos [91].

### Pangenome-wide inversions are enriched for transposable elements

Out of 755,656 insertions, a total of 54,313 contained DNA sequences with partial or complete sequence identity (>99%) with transposable elements (TEs). Sequence identity (>99%) with TEs was found in regions upstream (−100 bp) the breakpoints for 36,842 insertions, while 38,073 insertions had sequence identity with TEs downstream (+100 bp) of the breakpoints. For 1,166 inversions, 890 contained sequences with partial or complete sequence identity (>99%) with TEs. Total 136 inversions, had sequence identity with TEs in regions upstream (−100 bp) of the breakpoints, and 174 had sequence identity with TEs downstream (+100 bp) of the breakpoints. We identified 12 TE families. Insertions frequently contained LTR/Copia, RC/Helitron, CMC-EnSpM and LINE elements, while their breakpoints had a higher presence of LTR/Gypsy, LTR/Copia, and CMC-EnSpM elements. Sequences within inversions and their breakpoints exhibited a higher presence of LTR/Copia and LTR/Copia elements (Fig. 5).

**Figure 5:**
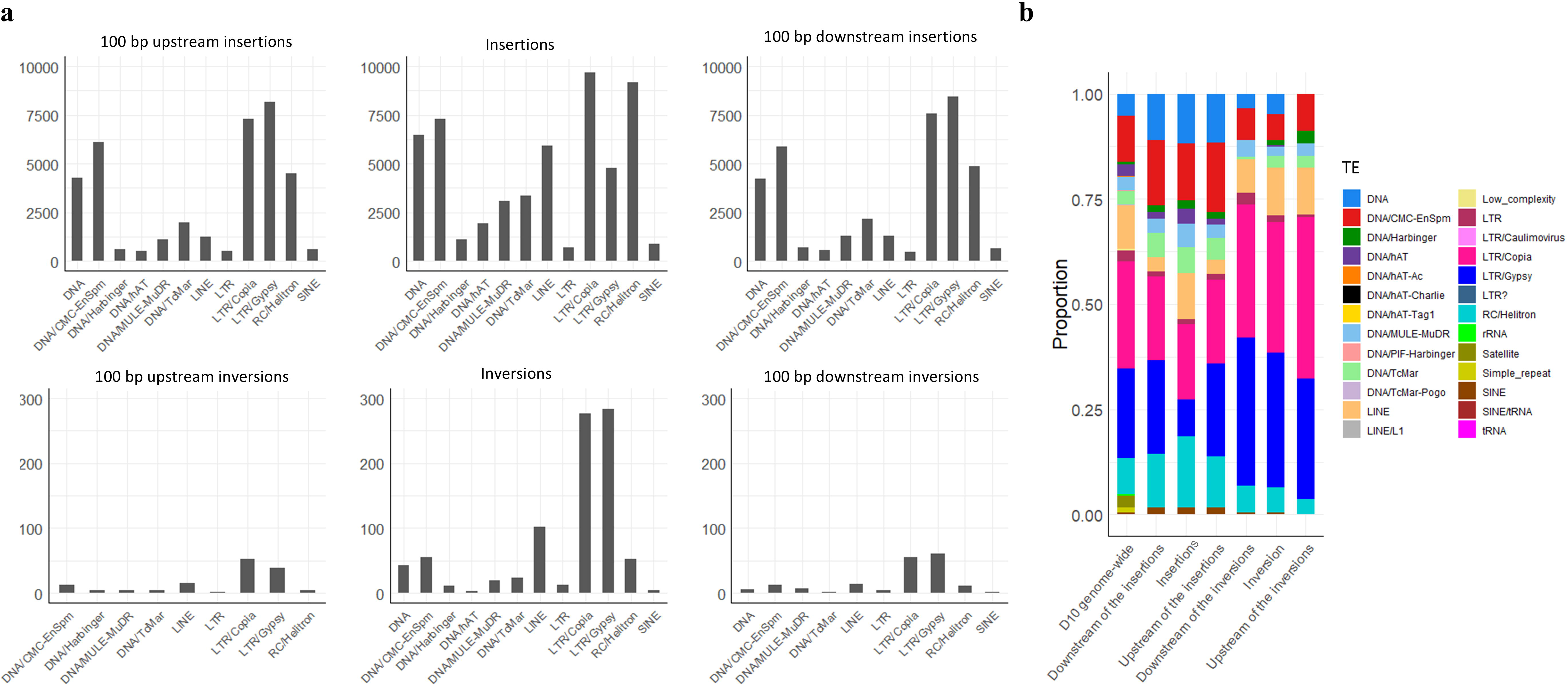
Overview of the distribution of transposable element (TE) families for insertion and inversion events in diverse accessions of *B. napus*. a) Frequencies (y-axis) of different TE families (x-axis) in pangenome-wide insertion (top) and inversion (bottom) events. Bar plots show the frequencies of TEs observed within SV, and in the 100 bp regions flanking SV breakpoints. b) Proportion (y-axis) of different TE families in pangenome-wide insertion and inversion events in comparison to Darmore v10 genome [40] (x-axis).

### Abundant repetitive sequences at breakpoints of collective SV and inversions

To investigate the mechanisms involved in the formation of insertions, deletions, and inversions, genomic regions flanking 100 bp upstream and downstream of putative breakpoints were searched for sequence motifs, as well as direct or inverted repeats. We separately analyzed insertion and deletion events from the collective SV. Deletions and insertions were divided into different size groups ranging from 30 bp to 30 kbp to assess whether different sequence motifs were associated with different size groups. This analysis revealed the presence of direct repeats (including microsatellites), and inverted repeats at the breakpoints of both ends for all investigated categories, suggesting a non-random distribution. The results showed significant enrichment of poly-A or poly-T stretches, and GC-rich palindromic sequences. Poly-A or poly-T sequences were abundant across events in all categories (Additional file 2: Fig. S1). For inversion events, different patterns of GC-rich palindromic sequences were found at the breakpoints (Additional file 2: Fig. S2).

## Discussion

In this study, we surveyed the SV landscape of 94 ecogeographically diverse *Brassica napus* accessions using long-read sequencing data and reference-guided genome assemblies. A *B. napus* pangenome SV Atlas was constructed describing distribution and frequencies for collective SV, inversions, and gene PAV, capturing species-wide insertions, deletions, inversions, and gene presence-absence variations across the divergent *B. napus* morphotypes. We found that the collective SV and inversions affect up to 15.14% and 11.47% of the genome, respectively. The pangenome-wide gene PAV indicated that 38.88% of *B. napus* genes were affected by large segmental chromosomal deletions in at least two accessions, and 78.97% by large segmental duplications. Using this diverse species-wide panel of accessions, our findings demonstrated that a broad spectrum of structural variant types is linked to intraspecific diversification in *B. napus*, emphasizing their collective significance in evolution and breeding of this important, recent allopolyploid crop species. Selective sweep signatures based on the distribution and frequency of collective SV and inversions, as well as gene PAV linked to intraspecific diversification, were predominantly located outside centromeric regions and in chromosome arms, which is consistent with the expected locations of chromosomal crossover [44] and the predominance of coding genes in *B. napus* [3]. Regions under selective pressure for intraspecific morphotype diversification contained putative genes associated with various adaptation mechanisms, such as flowering time, cell wall expansion, organ development, environmental response, abiotic stress response, and disease resistance, as previously described in studies exploring SNPs, InDels, CNVs, and homoeologous chromosomal exchanges across different *B. napus* ecotypes [3,13,21,26,28,29].

Our results showed a closer phylogenetic clustering of swede and East Asian OSR accessions to *B. rapa.* During the adaptation of rapeseed introduced from Europe to East Asia, repeated introgressions from Asian *B. rapa* into Asian *B. napus* cultivars have been documented [19,45]. Furthermore, evidence of interploidy gene flow from European turnip into swede has been reported [21]. It is accepted that *B. rapa* is the maternal progenitor of rapeseed [13]. Although the initial ancestral parents are unknown, the allopolyploid species is extremely recent compared to the 4.6 million years since *B. rapa* and *B. oleracea* diverged [46]. Nevertheless, introgressions from diploid progenitors and swede [23] during the adaptation history of OSR, along with the fact that no wild *B. napus* exists [13], makes it complicated to trace its order of divergence and ancestral lineage [22].

Our study employed insertions, deletions, inversions and gene presence-absence variations for a species-wide investigation of subgenome asymmetry, as well as an analysis of selective sweep signatures in *B. napus*. Previous studies utilized other types of genetic variations, such as SNPs [3,13,21,25,26], SSRs [27], CNVs [28], and homoeologous chromosomal exchanges [29]. Based on structural variations, we demonstrated an unbalanced distribution of SV, and an asymmetrical selection pattern between *B. napus* subgenomes. We observed a higher number of insertions and deletions in subgenome A compared to subgenome C across the panel. Furthermore, there were more genes with SV in coding sequences (CDS) and untranslated regions (UTRs) associated with R genes in subgenome A compared to C, while flowering-time-related genes were more prevalent in subgenome C than in subgenome A. The pangenome-wide distribution and frequency of collective SV revealed stronger selective sweep signatures in subgenome C compared to subgenome A when comparing swede vs. non-swede, and WOSR vs. SOSR, with the greatest difference observed for swede vs. non-swede. The observed asymmetry between the *B. napus* subgenomes is a complex phenomenon attributed to inherent characteristics of the progenitor species prior to hybridization, along with the influence of breeding procedures that utilized progenitors or closely related crops as donors. A previous study using a diversity panel representing variability in Chinese semi-winter rapeseed reported a higher rate of recombination and genetic diversity in subgenome A [47], likely reflecting past crossings between *B. rapa* and *B. napus* and the better viability of the resulting hybrid seeds compared to attempts to cross *B. oleracea* and *B. napus* [48,49]. Another study, which focused mainly on accessions from Asia, investigated divergence in gene function and differential selection pressures between the two subgenomes post-polyploidization, reporting stronger selective sweep signatures in subgenome C for (East Asian) semi-winter vs. winter, and spring vs. winter rapeseed. This study highlighted that during the breeding process from original *B. napus* to landraces, asymmetrical subgenomic selection of genes led to ecotype change. This included improvements in *B. napus* with respect to adaptation to environmental variation and oil accumulation through A subgenome-specific selection on defense-response genes and triglyceride biosynthetic genes, respectively [13].

Similarly, another study focusing on mostly Asian accessions reported more genes associated with plant defense and oil accumulation in subgenome A, whereas developmental-related genes, including flowering-time-related genes, were more prevalent in subgenome C compared to A [50]. Gene expression studies have also noted biases depending on the history of the plant material or tissue used. Some reported an expression bias toward subgenome A [51], while others observed a bias toward subgenome C [52].

When comparing swede vs. non-swede, East Asian OSR vs. all other OSR, and WOSR vs. SOSR accessions, our findings revealed that SV impacted genes with putatively roles in morphotype development, disease resistance and responses to environmental and abiotic stress. These included genes with SV-carrying variants coinciding with selective sweep signals or genes affected by large segmental presence-absence variations. For swede, the identified genes were putatively involved in storage organ formation, cell division and expansion, meiosis, and environmental responses and disease resistance, highlighting morphotypic differences between swedes and non-swedes. For East Asian OSR vs. all other OSR, regions under selection contained genes linked to environmental response, disease resistance, and abiotic stress response, aligning with historical records suggesting that rapeseed underwent breeding after its introduction from Europe to East Asia to adapt to new environments [13]. Similarly, in the WOSR vs. SOSR comparison, genes putatively involved in environment interaction, stress response, disease resistance, and flowering time were identified. The role of SV in differentiating WOSR and SOSR, particularly for flowering time genes such as *BnFLC.A03* and *BnFLC.A10*, has been well-documented [43,53]. Intragenic SV can affect gene variants differently depending on their location within coding and untranslated regions, as well as their size and number. However, the overlap of SV-carrying gene variants with selective signatures can imply their importance for further studies. Large chromosomal segment duplications and deletions also distinguished swede from OSR, particularly in subgenome C. For instance, gene presence-absence variation analysis found genes that are reported to have higher expression in swede compared to rapeseed during swollen hypocotyl formation [29], such as homoeologous gene pair *BnBLH1.A04 and BnBLH1.C04* which were located in a duplicated and deleted segment respectively. Results also revealed two copies of *BnFLC.C09* and *BnATX2.C08*, both putative flowering time inhibitor genes, and one copy of *BnCEL2.C08*, a putative cell wall development and expansion gene, in a deleted segment. Meanwhile, one copy each of *BnFLC.A10* and *BnATX2.A09*, and two copies of *BnCEL2.A09* were found within duplicated segments, offering a more detailed perspective compared to previous studies focused solely on flowering time genes [28].

Structural variant (SV) calling methods based on read alignment [54] allow for the filtering of SV by read depth, which enhances the accuracy of SV detection. However, despite advancements in long-read sequencing technology which have increased read lengths, this approach becomes less efficient for detecting large SV that exceed the read length. To address this limitation, we used a reference-guided genome assembly approach to detect inversions larger than 1 Mb. Nonetheless, this approach is still limited in detecting more complex SV, such as translocations or very large inversions, as the alignment procedure assumes a linear and collinear relationship with the reference genome.

We identified inversion hotspots within or near centromeric regions throughout the *B. napus* panel, consistent with reports that inversions in the Arabideae tribe are often enriched in these regions [55]. Such inversions can contribute to the suppression of meiotic crossovers, preserving the structural integrity of the centromeres and maintaining their proper function during cell division [55,56], which reduces the influence of selection in these regions [57].

Analysis of transposable element (TE) content in inversions showed an enrichment to 76.33% compared to the average TE content of 54% observed in the Darmor v10 genome [40]. Long terminal repeat (LTR) Copia and Gypsy elements were the most abundant TE families in inversion sequences and inversion breakpoints. Similar findings have been reported in a pangenome inversion index constructed for rice [58]. Differences associated with various TE families has been reported in the sequence of insertions located in similar position in different genotypes, indicating connection with diversification of *B. napus* [36]. Therefore, we considered both position and similarity of sequences, when merging SV from accessions throughout the panel. We investigated abundance of TE content in insertions which showed 7.2% of insertions had high sequence similarity to various TE families.

We investigated inversions and their connection to intraspecific diversification, and identified large inversions that are prominent in swedes and SOSR. Our findings showed that the strength and frequency of selective sweep signatures based on inversions increased, on average, outside centromeric regions and towards paracentric regions in all pairwise comparisons. This aligns with the reported role of inversions in species divergence and adaptation, by potentially preserving beneficial gene combinations while reducing the probability of crossover within the inverted region. Large inversions and large PAV are known to influence occurrence of recombination [59].

Large segmental chromosomal PAV and inversions distinguished swedes from non-swede accessions. For example, segmental deletions and duplications larger than 10 kbp in chromosomes A09, C01, C02, C08, and C09 differentiated swedes from all OSR ecotypes. Also, a deletion was found in all accessions of East Asian OSR in chromosomes C03 and another deletion in all WOSR accessions in chromosomes C05. These duplications or deletions may have arisen from unbalanced gametes formed due to meiotic recombination in paracentric regions [60], leading to non-reciprocal chromosomal exchanges. Individual swede accessions were reported earlier to carry a higher level of homoeologous chromosomal exchanges compared to OSR, particularly between choromosomes C01-A01, C08-A09, and C09-A10, and mostly in the direction of A subgenome segments replacing those of C [29]. This could be detected as deleted segments in subgenome C chromosomes such as C01 and C08 [3]. By demonstrating that these and other large SV are specific to particular crop morphotypes or ecotypes, we show that such variants were likely a key factor in the formation of novel phenological and physiological variation that was selected by humans during evolution of *B. napus* crops.

## Conclusion

In conclusion, our study provides a comprehensive survey of the pangenome-wide structural variations in the important crop *B. napus* and demonstrates that a broad spectrum of SV is associated with interspecific morphotype and ecotype diversification. The study provides valuable insights into SV associated with selection signals throughout the evolution and breeding of *B. napus*, as well as the underlying genes and genetic elements in affected chromosome regions. This comprehensive analysis of the SV landscape offers a valuable resource for further improving *B. napus* as a crop, for example by implementing SV in breeding. Additionally, larger SV, such as inversions and segmental deletions and duplications, can shed light on how these variants have influenced patterns of genomic recombination within each ecotype or subspecies, and help overcome chromosome regions which are recalcitrant for recombination.

## Methods

### Plant material

Aiming for species-wide detection of structural variations in *B. napus*, we selected 94 genetically and ecogeographically diverse accessions broadly representing the genetic diversity present in the ERANET-ASSYST diversity set, a species-wide panel of over 500 accessions previously described by Bus et al. [27]. The selected accessions originated primarily from Europe and East Asia, the main centers of origin for modern *B. napus* crop types, along with key oilseed and swede cultivars bred in North America and Australasia. Prior to sequencing, all accessions were maintained through controlled self-pollination for at least four generations in Plant Breeding Department at Justus Liebig University Giessen, Germany. Ecotype classification followed Bus et al. and included WOSR (22 accessions), SOSR (23 accessions), East Asian OSR (22 accessions), and swede/rutabaga (17 accessions), along with Asian *B. napus* vegetable forms and fodder rape accessions [27]. Detailed information about morphotypes, ecotypes and geographic origins for each accession is presented in Additional file 1: Table S1. Long-read sequencing data for two accessions of *B. rapa* (*B. rapa* ssp. *rapa* and *B. rapa* ssp. *broccolieto*) [61], and two accessions of *B. oleracea* (Genebank/cultivar IDs HRIGRU007343 and HRIGRU007338) [62], as well as genome assemblies of *B. rapa* Z1 (yellow sarson) and *B. oleracea* HDEM (broccoli) [63], were downloaded from public resources to represent the diploid progenitors.

### High molecular weight DNA isolation and long-read sequencing

Three plants from each accession were grown in a greenhouse to generate tissue. Fully expanded young leaves were collected, immediately deep-frozen in liquid nitrogen, and stored at −80°C for DNA extraction. High molecular weight (HMW) genomic DNA was extracted from 2.5 grams of the two youngest leaves from a single plant of each accession using the NucleoBond HMW DNA extraction kit (Macherey-Nagel, Germany). The kit protocols were followed according to the manufacturer’s instructions. DNA quality was assessed via 0.5% agarose gel electrophoresis (80V, 0.5×TBE, 60 min), a UV/Vis spectrophotometer (NanoDrop Technologies, USA), and a Qubit fluorometer using the Qubit dsDNA BR assay kit (Life Technologies, USA). Long-read DNA sequencing using Oxford Nanopore Technology was conducted by the DFG NGS Competence Center Tübingen (NCCT, Tübingen, Germany). HMW DNA was further qualified using the FP-1002 Genomic DNA 165-kbp kit for FEMTO Pulse systems (Agilent Technologies, USA). Library preparation was performed using the native ligation sequencing kit (SQK-LSK109; Oxford Nanopore Technologies), followed by sequencing using Oxford Nanopore Technology PromethION R9.4.1 flow cells. DNA fragments smaller than 25 kbp were progressively depleted before library preparation using the Short Read Eliminator kit (Circulomics, USA) to enhance read length and sequencing output. Base calling was performed using Guppy Basecaller v.5.1.12 and 6.1.5 (Oxford Nanopore Technologies) with the dna_r9.4.1_450bps_hac_prom model.

### Constructing reference-guided genome assemblies

*De novo* contig assembly was performed using the filtered long reads as input with Flye v2.9-b1768 [64]. The *de novo* assembled contigs were scaffolded based on the Darmor-bzh v10 (D10) reference genome [40] using Ragtag v2.1.0 [65]. N50, assembly size, and BUSCO scores were measured, and assembly statistics were calculated with Assembly-stats v1.0.1 [66]. The completeness of the assemblies was evaluated using BUSCO v5.7.1 [67] with the embryophyte_odb10 lineage dataset.

### Aligning long reads to reference genome and determining pangenome-wide distribution and frequency of collective SV

To remove any artificial bias from the raw long reads, they were filtered using NanoFilt v2.8.0 [68] (parameters: -l 5000 -q 10 --headcrop 60 --tailcrop 60). The quality of the filtered data was assessed using NanoPlot v1.40.0 [68], and genome-wide coverage was estimated. The trimmed raw reads were aligned to the D10 reference genome [40] using Minimap2 v2.24-r1122 [69] (parameters: -x map-ont --MD -a). The aligned reads were further filtered using the view function from Samtools v1.10 [70] (parameters: -bS -F 4 -h -q 50).

Structural variant (SV) calling was based on the methodology previously validated by Chawla et al. and Yildiz et al. [42,71], with addition of modifications that included an elaborate pipeline comprising multiple SV callers and rigorous filtering steps. Filtered aligned reads were used as input for SV calling with Sniffles v2.0.6 [72] (parameters: --minsupport $SV_minsupport -- minsvlen 30) and CuteSV v1.0.13 [73] (parameters: -s $SV_minsupport --genotype -l 30 -- max_cluster_bias_INS 100 --diff_ratio_merging_INS 0.3 --max_cluster_bias_DEL 100 -- diff_ratio_merging_DEL 0.3 --retain_work_dir --report_readid). The parameter “$SV_minsupport” was defined as 40% of the sequencing data coverage. Homozygous SV loci (allele frequency (AF) ≥0.9) with precise breakpoints were retained for each SV caller, removing SV detected in potentially heterozygous regions, which could indicate misaligned homoeologous segments.

To reduce the likelihood of false positives, for each accession the filtered call sets from CuteSV and Sniffles were merged using Survivor v1.0.7 [74] (parameters: 1000 2 1 1 −1 −1), retaining homozygous SV loci with precise breakpoints detected by both callers. These filtered SV were then used for a re-genotyping (force-calling) step using Sniffles, followed by additional filtering of the re-genotyped SV (parameters: PASS, QUAL ≥ 50, GQ ≥ 50, DV ≥ 12).

Due to the complexity of the *B. napus* genome, particularly the presence of homoeologous genomic regions and potential exchanges between subgenomes A and C, several scenarios were considered during the filtering of the SV call sets, including reciprocal and non-reciprocal chromosomal homoeologous exchanges. Notably, non-reciprocal homoeologous exchanges, where two copies of a chromosomal segment from one subgenome exist while the corresponding segment in the other subgenome is deleted, can be detected as duplicated and deleted segments by analyzing the coverage of long reads aligned to the reference genome. A read coverage calculation pipeline detailed by Stein et al. [75] used the aligned bam files to estimate the read coverage, and the areas with lower read coverage are considered as deleted chromosomal segments. To filter-out the SV that overlapped these segments (≥10 kbp), the following procedure was applied using this pipeline with modification: Aligned reads were used to calculate coverage across chromosomes using the bamtobed and genomecov functions from BEDTools v2.30.0 [76], which served as input for the modified deletion-duplication pipeline [75]. Outlier regions with coverage above 150 were discarded, and segments ≥10 kbp with coverage deviating by at least one standard deviation below the mean were considered as deleted segments. The resulting BED file was used as input for the intersect function from BEDTools to remove SV from the call set that overlapped with large deleted segments.

Finally, SV from all accessions were merged using Jasmine v1.1.5 [77] (parameters: max_dist=1000 min_support=1 min_seq_id=0.9 --nonlinear_dist --output_genotypes -- normalize_type). This software allows to account for sequence identity to differentiate SV based on both position and sequence similarity. For the merging process, variants within 1000 bp of each other were considered the same, provided they shared at least 90% sequence identity, and single-sample variants were also retained. For validation, SV were further visualized with the Integrative Genomics Viewer (IGV) tool [78] and compared to previously published SV. Long-read sequencing data for *B. rapa* and *B. oleracea* were aligned to their respective D10 subgenomes, A and C.

### Annotation of putative functional SV

To annotate putative functional SV, the intersect function from BEDTools v2.30.0 [76] was used to identify overlaps between SV events and gene models in the GFF file of the D10 reference genome. Araport11 *A. thaliana* representative gene model complementary DNA (cDNA) sequences [79] from TAIR were downloaded to create a local database using BLASTn [80]. Functional annotation of genes was carried out by aligning the cDNA sequences of the D10 reference genome against the local database (parameters: -evalue 1e-4 -perc_identity 90 - ungapped -max_target_seqs 1 -max_hsps 1). A list of *Arabidopsis* flowering time regulators from the Flowering Interactive Database (http://www.phytosystems.ulg.ac.be/florid) [81] and a list of putative R genes for *B. napus* published by Dolatabadian et al. [82] were used to facilitate gene function annotation.

### Detecting inversions using reference-guided genome assemblies and determining pangenome-wide distribution and frequency

Inversions larger than 30 bp were detected using the 94 reference-guided genome assemblies following the approach detailed by Zhou et al. [58], with modifications, including the use of Minimap2 as the aligner [83]. In summary, each genome assembly was aligned to the D10 reference genome using Minimap2 (v2.15) (parameters: -ax asm5 --eqx --cs -c --MD). Regions of synteny and inversions were identified using SyRI v1.6.3 [54], and inversions were merged using Jasmine v1.1.5 [77], with filtering applied to the aligned sequences for a minimum identity of 90% (parameters: max_dist=1000 min_support=1 min_seq_id=0.9 --nonlinear_dist -- output_genotypes --normalize_type). SV events marked as <INV> and with a “PASS” status were retained. Inversions correlating with ecotype diversification for WOSR, SOSR, East Asian OSR and swede/rutabaga accessions were separated into subsets based on the following criteria: ecotype of interest ≥70%, other ecotypes ≤30%. East Asian OSR accessions were not included as “other ecotypes” because some accessions in this group clustered closely with WOSR and SOSR accessions in principal component and phylogenetic analyses. The under-represented inversions were determined by ecotype of interest ≤70%, other ecotypes ≥30%. The filtering was performed in R [84], therefore limitations in floating-point precision might occur. The assemblies of *B. rapa* and *B. oleracea* was aligned to their respective D10 subgenomes, A and C.

### Chromosomal distribution of pangenome SV hotspots

For an overview of distribution and localization of SV hotspots, we performed a 200 kbp fixed-window analysis across all chromosomes using the coverage function of BEDTools. The top 30% of windows with the highest frequency of inversion start coordinates were defined as hotspots, separately for each subgenome. To compare subgenomes A and C, number of events per each 200 kbp were standardized by percentage for whole genome and compared using one-tail Student’s test (α=0.01).

### Phylogenetic analysis and principal component analysis

For phylogenetic analysis, the pipeline was first tested using the merged SNP call set of all accessions. Then, the merged SV events for all accessions were used to construct maximum likelihood (ML) trees using IQ-TREE v2.2.0.3 [85], based on the best model determined by the Bayesian Information Criterion (BIC). Two accessions of *B. rapa* (*B. rapa* ssp. *Rapa* and *B. rapa* ssp. *broccolieto*) [61], and two accessions of *B. oleracea* (Genebank/cultivar IDs HRIGRU007343 and HRIGRU007338) [62] as well as genome assemblies of *B. rapa* Z1 (yellow sarson) and *B. oleracea* HDEM (broccoli) [63], were used to represent diploid progenitors. The analysis was performed separately for the A and C lineages based on the D10 reference genome. For collective SV, 25% of each chromosome end was excluded to avoid using regions prone to homoeologous recombination, and the software selected the BIC model GTR2+FO+G4. The reliability of the ML trees was estimated using the ultrafast bootstrap approach (UFboot) with 1000 replicates, and the consensus ML trees were visualized using the online tool Interactive Tree of Life (iTOL) v6.8 (https://itol.embl.de). The same procedure for phylogenetic analysis was applied to the inversions, but no chromosomal segments were excluded. The software selected the BIC model GTR2+FO+ASC+G4. Principal component analysis (PCA) of the merged SV events was performed for the whole genome using Plink v1.90b6.21 [86], and visualized with the plotly package v4.10.1 in R [84].

### Characterization of selective sweep signals

To identify potential genomic regions under selection, the pairwise fixation index (FST; Weir and Cockerham FST estimates) [87] was calculated for pairwise comparisons of swede vs. non-swede, East Asian OSR vs. all other OSR, and WOSR vs. SOSR accessions using the VCFtools suite [88] across non-overlapping 50 kbp windows. FST values were standardized and transformed into z-scores in R [84] using the formula: Z = (x - μ) / σ, where x is the FST value for each bin, μ is the mean of FST values, and σ is the standard deviation of FST values. Genomic regions under selection were identified using a one-tailed Z-test with a significance level of α = 0.05, corresponding to Z = 1.645. Using the intersect function of BEDTools, the results were intersected with the gff file of the D10 reference genome to find genes located in regions under selective signatures, and then merged with annotations from Araport11 using R [84]. For each pairwise comparison, the genetic loci that have z values >0 were used to compare the average z values among subgenomes using one-tail Student’s test with a significance level of α=0.01. The GO enrichment analysis was performed using goseq package [89] in R, with a significance level of α = 0.05.

### Detecting large duplicated or deleted genomic segments and determining pangenome-wide gene PAV

To detect genomic rearrangements, aligned reads were used to calculate coverage across chromosomes using the bamtobed and genomecov functions from BEDTools, which served as input for a modified deletion-duplication pipeline based on Stein et al. [75]. Briefly, outlier regions with coverage above 150 were discarded, and segments ≥10 kbp with coverage deviating by at least one standard deviation above or below the mean were identified as duplications and deletions, respectively. Using the intersect function from BEDTools, the results were compared with the gff file of the D10 reference genome to detect genes affected by large chromosomal segments that were either deleted or duplicated.

To identify genes whose presence in these deleted or duplicated regions correlated with ecotype diversification among WOSR, SOSR, East Asian OSR, and swede accessions, the following criteria were applied: ecotype of interest ≥70%, other ecotypes ≤30%. The filtering was performed in R [84]. The results were then merged with Araport11 gene functional annotations using R [84]. East Asian OSR accessions were not included as “other ecotypes,” as some accessions from this group clustered closely with WOSR and SOSR accessions in principal component and phylogenetic analyses.

### Annotation and analysis of TEs

To annotate the potential overlap of TEs with insertions and inversions, the *Brassica napus* transposable element library, first described by Chalhoub et al. [3], was downloaded from the supplementary files provided by Rousseau-Gueutin et al. [40] and used to create a local database with BLASTn [80]. Then, in two separate procedures for insertions and inversions, their sequence and the sequences 100 bp upstream and downstream of their putative breakpoints were blasted against the local database (parameters: -evalue 1e-4 -perc_identity 99 -ungapped - max_target_seqs 1 -max_hsps 1).

### Characterization of sequence motifs in SV breakpoints

For characterizing the SV-breakpoints of merged SV across the panel, sequences 100 bp upstream and downstream of each putative SV-breakpoint was extracted. The analysis was performed using the motif finder MEME v5.4.1 [90] as described by Samans et al. [4]. For the collective SV, merged SV were grouped into seven different size ranges: 30-500 bp, 501-1000 bp, 1001-2000 bp, 2001-3000 bp, 3001-10000 bp, 100001-30000 bp, 30001-100000 bp to account for the possibility of different SV-origin mechanisms.

### Availability of data and materials

All raw data generated in this study including Long-read DNA sequence data are available on the NCBI BioProject database (https://www.ncbi.nlm.nih.gov/bioproject/) under accession number PRJNA1183293. Genome assemblies constructed in this study are available on the European Nucleotide Archive (https://www.ebi.ac.uk/ena/browser/home) under accession number PRJEB82438.

## Acknowledgement

NGS methods including preparing NGS libraries and generating Oxford Nanopore sequencing data were arranged and performed with the support of Elena Buena Atienza, Caspar Gross, and Nicolas Casadei from NGS Competence Center Tübingen (INST 37/1049-1), funded by the Deutsche Forschungsgemeinschaft (DFG, German Research Foundation); No. of DFG proposal: SEQ2013, Project Number: 21103. We acknowledge provision of computing resources by Justus Liebig University Bioinformatics Core Facility (BCF), and de.NBI Cloud within the German Network for Bioinformatics Infrastructure (de.NBI) and ELIXIR-DE (Forschungszentrum Jülich and W-de.NBI-001, W-de.NBI-004, W-de.NBI-008, W-de.NBI-010, W-de.NBI-013, W-de.NBI-014, W-de.NBI-016, W-de.NBI-022). We also sincerely thank Sabine Frei and Regina Illgner for technical support with the greenhouse and lab experiments.

## Authors’ contributions

NPA was involved in conceptualization, methodology, designing and performing the experiments, data generation and analysis, validation, visualization, interpretation, writing the original draft of the manuscript; review & editing of the manuscript. AAG was involved in conceptualization, methodology, review & editing of the manuscript. RJS was involved in conceptualization, methodology, funding acquisition, and review & editing of the manuscript. All the authors have read and approved the final manuscript.

## Ethics declarations

### Ethics approval and consent to participate

Not applicable.

### Consent for publication

Not applicable.

## Supplementary Information

**Additional file 1: Supplementary Tables.**

Supplementary Table 1. List of 94 ecogeographically diverse rapeseed (*Brassica napus* L.) accessions selected from the ERANET-ASSYST diversity set.

Supplementary Table 2. Summary of long-read DNA resequencing data by Oxford Nanopore Technology for 94 *B. napus* accessions.

Supplementary Table 3. Genome assembly statistics for 94 *B. napus* accessions.

Supplementary Table 4. Number of removed SV in each chromosome in the process of filtering-out SV that are likely to overlap with a deleted chromosome segment.

Supplementary Table 5. Number of removed SV in the process of filtering-out SV that are likely to overlap a deleted chromosome segment.

Supplementary Table 6. Number of SV called for each *B. napus* accession by aligning long-reads to Darmor-bzh v10 reference genome.

Supplementary Table 7. Number of total SV called for each subgenome of *B. napus* accession by aligning long reads to Darmor-bzh v10 reference genome.

Supplementary Table 8. Overview of pangenome-wide distribution and frequency of collective SV in *B. napus*.

Supplementary Table 9. Overview of genes intersecting pangenome-wide collective SV in *B. napus*.

Supplementary Table 10. Overview of putative flowering-time-related genes intersecting pangenome-wide collective SV in *B. napus*.

Supplementary Table11. Overview of putative R genes intersecting pangenome-wide collective SV in *B. napus*.

Supplementary Table 12. Number of inversions called for each *B. napus* accession by aligning reference-guided genome assemblies to Darmor-bzh v10 reference genome.

Supplementary Table 13. Overview pangenome-wide distribution and frequency of inversions in *B. napus*.

Supplementary Table 14. Overview of distribution and frequency of over-represented or under-represented inversion events across *B. napus* pangenome for different ecotype groups.

Supplementary Table 15. Overview of collective SV hotspots across *B. napus* pangenome.

Supplementary Table 16. Overview of inversion hotspots across *B. napus* pangenome.

Supplementary Table 17. Fst Values for pairwise comparison of swede versus non-swede accessions using pangenome-wide collective SV in *B. napus*.

Supplementary Table 18. Fst Values for pairwise comparison of East Asian OSR versus all other OSR accessions using pangenome-wide collective SV.

Supplementary Table 19. Fst Values for pairwise comparison of WOSR vesus SOSR accessions using pangenome-wide collective SV.

Supplementary Table 20. Overview of putative genes intersecting genomic regions under selection for pairwise comparison of swede versus non-swede accessions using pangenome-wide collective SV.

Supplementary Table 21. Overview of putative genes intersecting genomic regions under selection for pairwise comparison of Asian OSR versus all other OSR accessions using pangenome-wide collective SV.

Supplementary Table 22. Overview of putative genes intersecting genomic regions under selection for pairwise comparison of WOSR versus SOSR accessions using pangenome-wide collective SV.

Supplementary Table 23. GO enrichment analysis for genes overlapping selective sweeps for pairwise comparison of swede versus non-swede accessions for pangenome-wide collective SV.

Supplementary Table 24. GO enrichment analysis for genes overlapping selective sweeps for pairwise comparison of East Asian OSR versus all other OSR accessions for pangenome-wide collective SV.

Supplementary Table 25. GO enrichment analysis for genes overlapping selective sweeps for pairwise comparison of WOSR versus SOSR accessions for pangenome-wide collective SV.

Supplementary Table 26. Fst Values for pairwise comparison of swede versus non-swede accessions using pangenome-wide inversions in *B. napus*.

Supplementary Table 27. Fst Values for pairwise comparison of Asian OSR versus all other OSR accessions using pangenome-wide inversions.

Supplementary Table 28. Fst Values for pairwise comparison of WOSR versus SOSR accessions using pangenome-wide inversions.

Supplementary Table 29. Overview of putative genes intersecting genomic regions under selection for pairwise comparison of swede versus non-swede accessions using pangenome-wide inversions.

Supplementary Table 30. Overview of putative genes intersecting genomic regions under selection for pairwise comparison of Asian OSR versus all other OSR accessions using pangenome-wide inversions.

Supplementary Table 31. Overview of putative genes intersecting genomic regions under selection for pairwise comparison of WOSR versus SOSR accessions using pangenome-wide inversions.

Supplementary Table 32. GO enrichment analysis for genes overlapping selective sweeps for pairwise comparison of swede versus non-swede accessions for pangenome-wide inversions.

Supplementary Table 33. GO enrichment analysis for genes overlapping selective sweeps for pairwise comparison of East Asian OSR versus all other OSR accessions for pangenome-wide inversions.

Supplementary Table 34. GO enrichment analysis for genes overlapping selective sweeps for pairwise comparison of WOSR versus SOSR accessions for pangenome-wide inversions.

Supplementary Table 35. Overlapping regions of pangenome-wide collective SV and inversions under selective sweeps in swede versus non-swede comparison.

Supplementary Table 36. Overlapping regions of pangenome-wide collective SV and inversions under selective sweeps in Asian OSR versus all other OSR comparison.

Supplementary Table 37. Overlapping regions of pangenome-wide collective SV and inversions under selective sweeps in WOSR versus SOSR comparison.

Supplementary Table 38. Overview of pangenome-wide distribution and frequency of gene presence-absence variation in *B. napus*.

Supplementary Table 39. Overview of pangenome-wide gene presence-absence that correlate with WOSR, SOSR, East Asian OSR, or swede group of accessions.

## Additional file 2: Supplementary Figures.

**Supplementary Figure 1:**
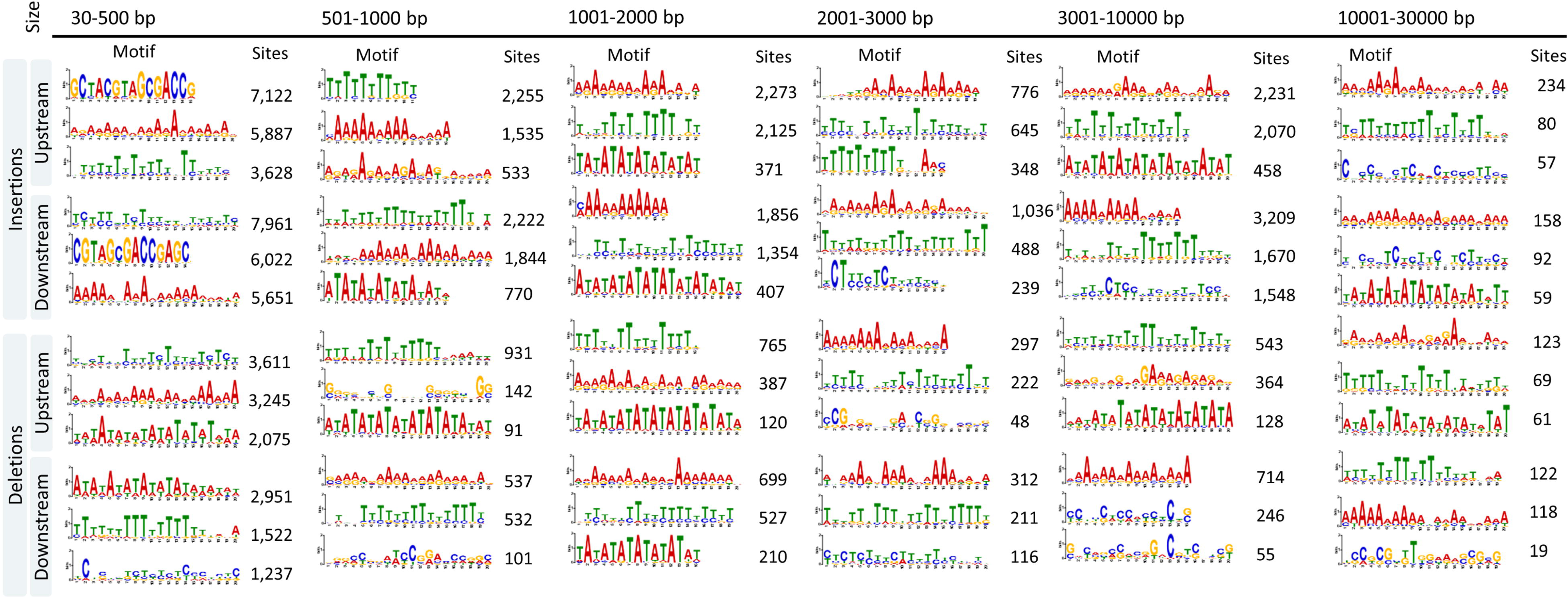
Motifs enriched around putative breakpoints of pangenome-wide insertion and deletion events in diverse accessions of *B. napus*. For each breakpoint, the genomic sequence of 100 bp upstream and downstream was extracted from six different size categories of SV and analyzed for enriched DNA motifs using MEME software, with accepted e-values < 0.05. For each SV size group, the three most abundant motifs (by the number of sites) are shown for both the downstream and upstream regions of the SV.

**Supplementary Figure 2:**
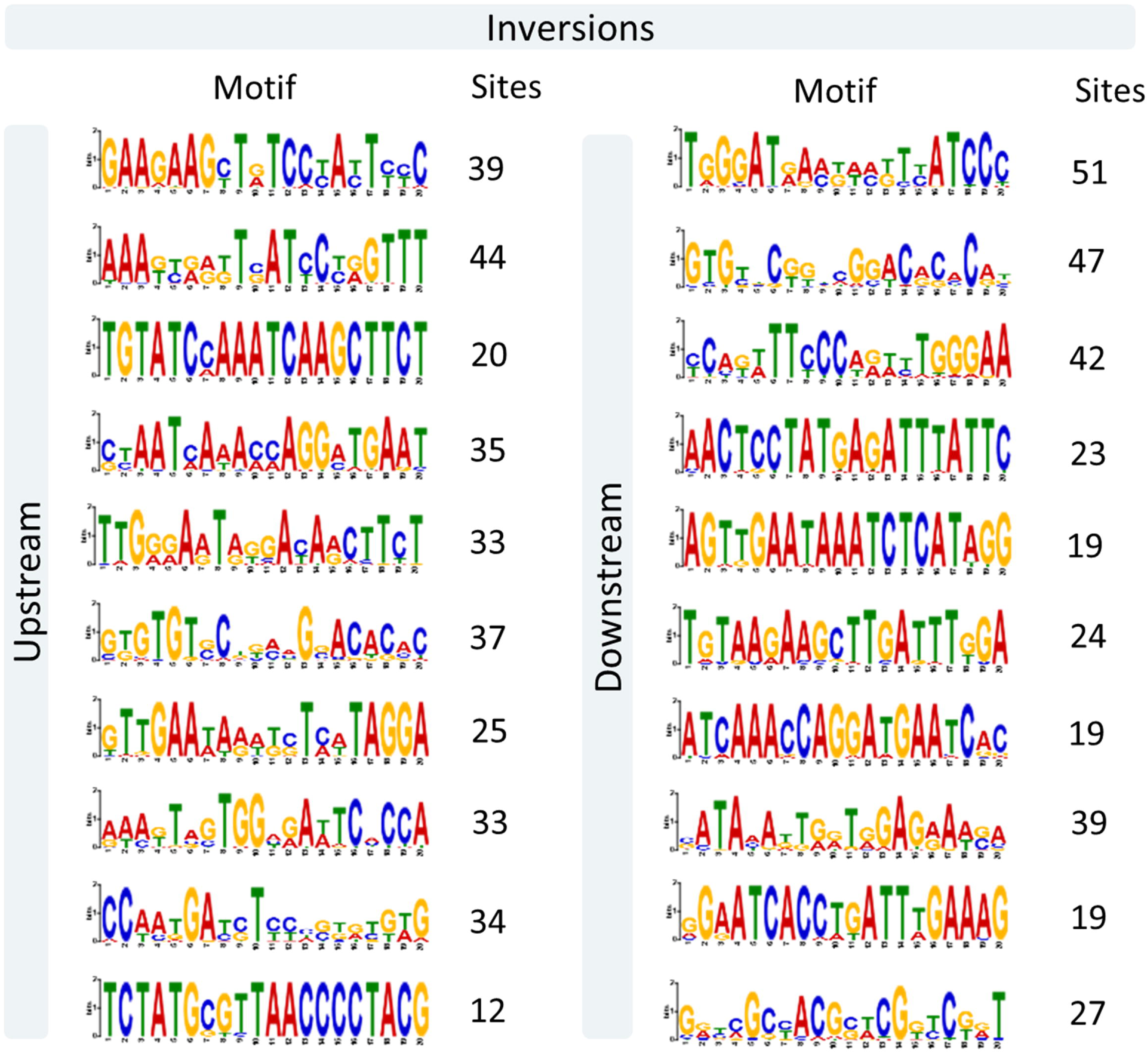
Motifs enriched around putative breakpoints of pangenome-wide inversion events in diverse accessions of *B. napus*. For each breakpoint, the genomic sequence of 100 bp upstream and downstream was extracted and analyzed for enriched DNA motifs using MEME software, with accepted e-values < 0.05.

